# Hepatocellular Carcinoma Revisited from Single Cell Sequencing: Dynamic Evolution from Epithelial Dedifferentiation to Mesenchymal Remodeling

**DOI:** 10.64898/2026.04.24.720317

**Authors:** Kai Yan, Wei Dong, Yeye Wu, Zhe Han, Jinling Hong, Hongbin Ma, Chengguang Zhu, Yangjie Xiong, Zhao Yang

## Abstract

**Background:** Single-cell RNA sequencing has provided new insights into hepatocellular carcinoma (HCC); however, a unified understanding of epithelial heterogeneity and immune evasion strategies in HCC remains lacking.

**Methods:** We re-analyzed publicly available single-cell datasets using conventional bioinformatics pipelines. Cell-type annotation of epithelial and T cell populations was further validated across multiple independent datasets to ensure robustness.

**Results:** We systematically examined epithelial, immune, myeloid, and stromal lineages. In addition to recapitulating previously reported findings, we identified several novel observations. Notably, we uncovered a three-step dedifferentiation trajectory in epithelial cells and confirmed a bidirectional differentiation pattern within CD8□ T cells. We also identified a subset of GZMK□ CD4□ T cells, whose transcriptional features resemble but are distinct from T follicular helper (Tfh) cells. Importantly, transcriptional drift within myeloid populations appeared to be closely associated with immune responsiveness. Furthermore, ligand–receptor analysis highlighted a potential cooperative role of LAMP3□ dendritic cells and Tfh cells in promoting lymphoid follicle formation.

**Conclusions:** In the era of rapidly evolving single-cell sequencing technologies, we provide a framework for understanding cellular heterogeneity in HCC, which awaits further validation in future studies.

## Introduction

Single-cell RNA sequencing has revealed pervasive heterogeneity in hepatocellular carcinoma (HCC), consistently highlighting tumor cell plasticity, T-cell exhaustion, and large-scale remodeling of the immune microenvironment^1-8^. However, these studies were typically conducted in separate cohorts and analytical frameworks, making it difficult to distinguish robust biological principles from cohort-specific or methodological biases. A systematic cross-dataset reanalysis is therefore needed to identify reproducible cellular programs and establish a unified view of the HCC ecosystem.

In this study, we revisited published single-cell datasets spanning malignant hepatocytes, immune populations, and stromal cells to uncover conserved features across cohorts. By paralleled study of multiple datasets using classical single cell analytic strategies, we aimed to identify recurrent transcriptional states and differentiation trajectories that are consistently observed despite technical and cohort heterogeneity.

Through this integrative analysis, we identify a conserved three-step dedifferentiation trajectory of malignant hepatocytes, a bidirectional differentiation program of CD8□ T cells associated with immunotherapy responsiveness, and a previously underappreciated cytotoxic CD4□ T-cell subset. Together, these findings provide a robust and reproducible single-cell view of the HCC tumor ecosystem and offer new insights into immune regulation and therapeutic response.

## Methods

### Data

The Lu dataset was used to analyze epithelial and stromal cells^4^. The Sharma dataset was used to validate epithelial cell annotations^7^. The Magen dataset was used to analyze immune cells^9^. Single cell study from Liu et al. was used to confirm the bidirectional differentiation of T cells in another organ^10^. Single-cell analysis data of normal liver were used as a reference and served as a benchmark for cell annotation^11-14^.

### Routine processing of single cell RNA sequencing data

All single-cell datasets were processed using a unified pipeline including SoupX for ambient RNA correction and scDblFinder for doublet removal^15,16^. Both tools were applied with default settings, as they do not require manual threshold tuning. Because raw empty-droplet matrices were unavailable for most datasets, the soup profile was estimated directly from the count matrix following the SoupX “no-empty-droplet” workflow. All single-cell datasets were further processed using standard workflows implemented in either Seurat (R) or Scanpy (Python), including normalization, highly variable gene selection, scaling, dimensionality reduction, and clustering^17,18^. Batch effects were corrected using Harmony (R) or scVI (Python), with inter-sample variability treated as a major source of batch effects^19,20^. For epithelial cells, clustering results obtained with Seurat were independently reproduced and validated using scVI to ensure robustness across analytical frameworks.

### Annotation and fraction analysis

Cell subpopulations were annotated based on upregulated genes and their functions, with reference to classic single-cell analysis literatures^11-13^. The analysis of cell subtype proportions in Python was performed using the scCODA package^21^. Metabolic pathway analysis was performed using the scMetabolism package^22^.

### Trajectory analysis

T-cell differentiation trajectories were reconstructed using Monocle3^23^. The UMAP embedding and cell annotations were directly inherited from the scVI clustering results. Trajectory graph learning was performed with customized parameters for each lineage. For CD4 T cells, *learn_graph* was run with *geodesic_distance_ratio* = 0.3, *euclidean_distance_ratio* = 1, and *minimal_branch_len* = 8. For CD8 T cells, parameters were set to *geodesic_distance_ratio* = 0.3, *euclidean_distance_ratio* = 0.1, and *minimal_branch_len* = 15. Changes in marker gene expression along pseudotime were analyzed following the strategy described by Ma et al^24^.

### Transcription factor analysis

For epithelial cells and T cells, the transcriptional drivers underlying observed differences were further investigated using pySCENIC^25^. The regulon specificity score (RSS) plot was used as a canonical approach to visualize transcription factor specificity across cell states.

### Public bioinformatics resources

Bulk RNA expression differences in the TCGA cohort were obtained using the GEPIA2 database^26^. DNA methylation data for TAL1 were retrieved from the UCSC Xena platform and compared between tumor and normal liver tissues^27^. Protein-level expression of selected marker genes was visualized using the Human Protein Atlas (HPA) database^28^.

## Results

### Single-Cell Dissection of Malignant Hepatocyte Dedifferentiation and Epigenetic Rewiring in HCC

We used the dataset generated by Lu et al. to analyze epithelial cells in hepatocellular carcinoma (HCC)^4^. This dataset spans multiple disease stages. Single-cell profiles were filtered using SoupX and scDblFinder^15,16^. We first focused on tumor heterogeneity. Epithelial cells originating from tumor tissues were directly regarded as malignant hepatocytes, without additional filtering using methods such as InferCNV, because normal hepatocytes are rarely present within highly malignant HCC lesions^29^. In total, 18,348 malignant hepatocytes (barcodes) were obtained. Unsupervised clustering separated these cells into three states: highly differentiated cells (high_diff_cell), dedifferentiated cells (de_diff_cell), and proliferating cells (prolif_cell), characterized by the markers HP/ARG1/CPS1, TFF2/S100A6, and STMN1/TOP2A, respectively (Fig. 1A, B). Across tumor progression (TNM stage I–IV), the proportion of dedifferentiated cells gradually increased (Fig. 1C). This “three-step” dedifferentiation and stemness-acquisition trajectory was validated in an independent dataset (Supplementary Fig. 1A, B)^7^. To exclude algorithm-driven bias, we reanalyzed the data using scVI and obtained consistent results (Supplementary Fig. 1C).

**Fig 1:**
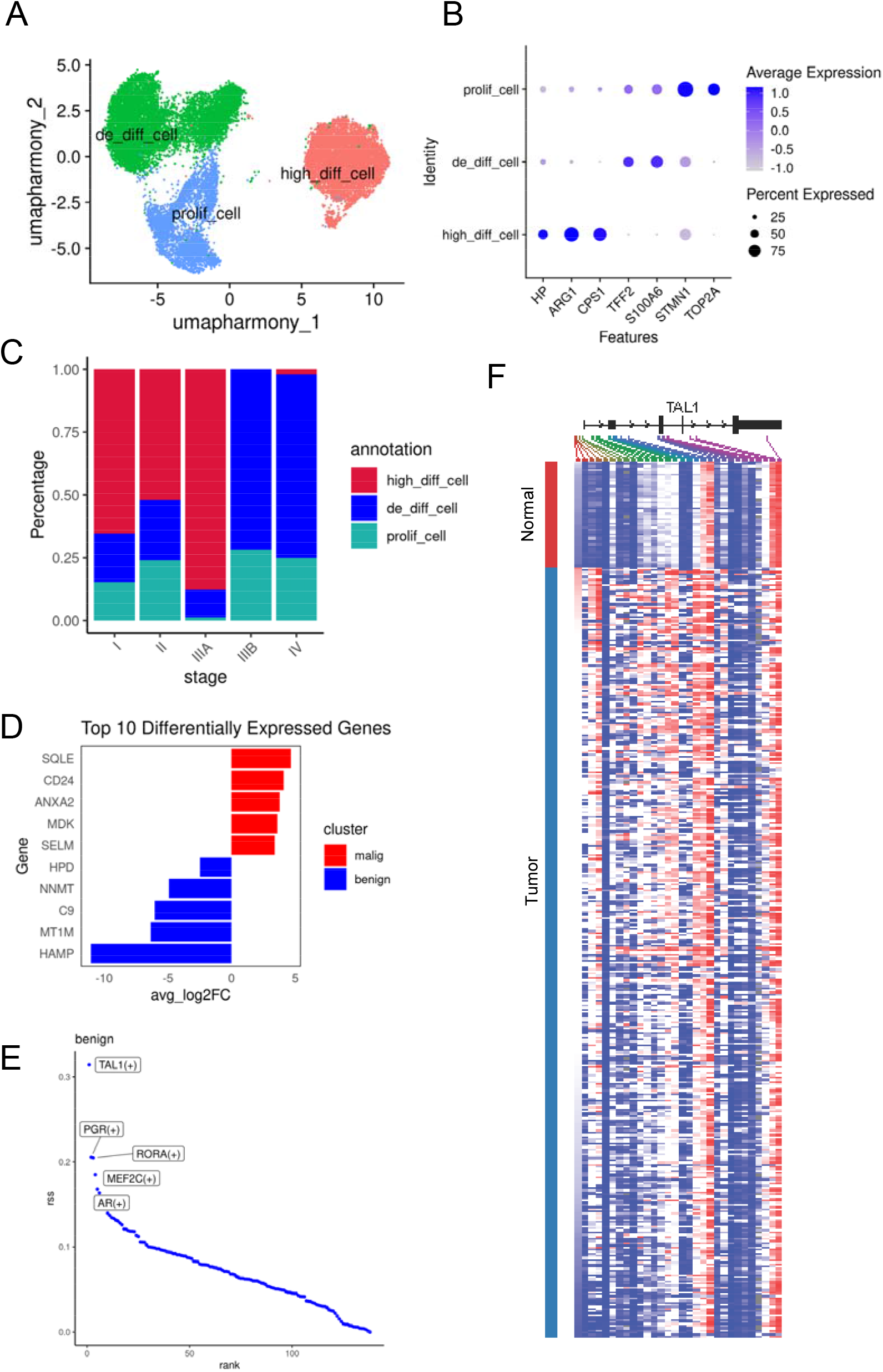
Heterogeneity of hepatocellular carcinoma cells and TAL1-associated regulatory alterations. (A) UMAP showing three cell states (de_diff_cell, high_diff_cell, prolif_cell) of hepatocellular carcinoma cells, with batch effects corrected using Harmony. (B) Dot plot of selected genes showing average expression (color intensity) and cell fraction (dot size) across the three states. (C) Stacked bar plot showing proportions of each cell state across AJCC stages. (D) Bar plot showing top 10 differentially expressed genes between malignant (red) and benign (blue) clusters. (E) Regulatory specificity score (RSS) plot for the benign liver cells regulatory program inferred by pySCENIC. (F) Comparison of TAL1 DNA methylation levels and CpG site signals between tumor and normal tissues (UCSC Xena).

We next compared the transcriptomes of malignant hepatocytes with normal hepatocytes (Fig. 1D). The top five upregulated genes in HCC were CD24, MDK, SELM, ANXA2, and SQLE, which are involved in immune evasion, cell proliferation, selenium metabolism, WNT signaling, and cholesterol metabolism, respectively. In contrast, normal hepatocytes showed higher expression of HPD, HAMP, NNMT, MT1M, and C9, which are associated with tyrosine metabolism, iron metabolism, xenobiotic metabolism, metal homeostasis, and humoral immunity, reflecting normal liver physiological functions. Notably, C9 is the terminal component of the complement cascade and forms the membrane attack complex (MAC) together with C5b–C8, directly inserting into tumor cell membranes to induce osmotic lysis^30,31^. Recent evidence further shows that GZMK can activate the complement pathway by cleaving C2 and C4^32^. GZMK□ CD8□ T cells are widely present in the tumor microenvironment; thus, downregulation of C9 in HCC may represent an important mechanism of immune evasion.

We then applied pySCENIC to identify transcription factors across differentiation states and obtained 138 regulons (Supplementary Fig. 2). Regulon specificity scores indicated that TAL1 is a key transcription factor associated with the normal hepatocyte phenotype (Fig. 1E). Analysis of the UCSC Xena database showed that TAL1 mRNA expression is not markedly altered in HCC; however, its DNA methylation level is significantly increased (Fig. 1F, Supplementary Fig. 3A). Elevated TAL1 methylation may result from aberrant activation of DNA methyltransferases (DNMTs)^33^. Consistently, public datasets show that DNMT3A and DNMT3B are significantly upregulated at the transcriptional level in HCC (Supplementary Fig. 3B).

### The Diverse Differentiation Potential of T Cells and Its Association with Immunotherapy Response

To investigate the heterogeneity and therapy-responsiveness of tumor-infiltrating T cells, we reanalyzed single-cell RNA-sequencing data from the Magen cohort^9^. This dataset includes samples with annotated PD-1 therapy response information, making it particularly suitable for dissecting the cellular and transcriptional features associated with resistance or sensitivity to immune checkpoint blockade. In addition, this dataset contains a large number of cells, enabling more robust and high-resolution clustering of immune populations (Supplementary Fig. 4A, B).

Single-cell transcriptomic profiling revealed extensive heterogeneity within tumor-infiltrating T cells, encompassing conventional CD4 □ and CD8 □ subsets, MAIT, NKT, and a minor γδ-like population (Supplementary Fig. 4C-E). While canonical CD4 □ and CD8 □ subsets were readily identified, several transcriptionally distinct subpopulations emerged, reflecting adaptive responses to the tumor microenvironment (TME). Notably, GZMB □ cells included both CD8 □ and CD4 □ phenotypes, suggesting a shared cytotoxic transcriptional program (Supplementary Fig. 4D, E)^34^.

Among CD4 □ T cells, trajectory analysis delineated multiple differentiation branches, including regulatory, follicular helper (Tfh), and two cytotoxic-like lineages (GZMK and GZMB) (Fig. 2A, B). A notable discovery was a GZMK □ CD4 subcluster co-expressing PDCD1, which was preferentially enriched in PD-1 therapy responders (Fig. 2C, D). These cells do not express the canonical Tfh markers CXCL13 and CD200 and therefore do not belong to the classical Tfh lineage (Supplementary Fig. 5A). This subset likely represents an active, PD-1–responsive helper population bridging canonical Tfh and cytotoxic states^35^. We also observed the presence of GZMB□ CD4 T cells. This population was also confirmed in another dataset (Supplementary Fig. 5B)^10^. Since GZMB□ CD4 T cells are enriched in normal liver tissue, we speculate that they may represent a liver-specific immune niche (Fig. 2E). An association between the Tfh population and responsiveness to PD-1–based immunotherapy was also observed (Fig 2D, Supplementary Fig. 5C). Molecular expression changes accompanying CD4 T-cell differentiation were also characterized (Supplementary Fig. 6).

**Fig. 2:**
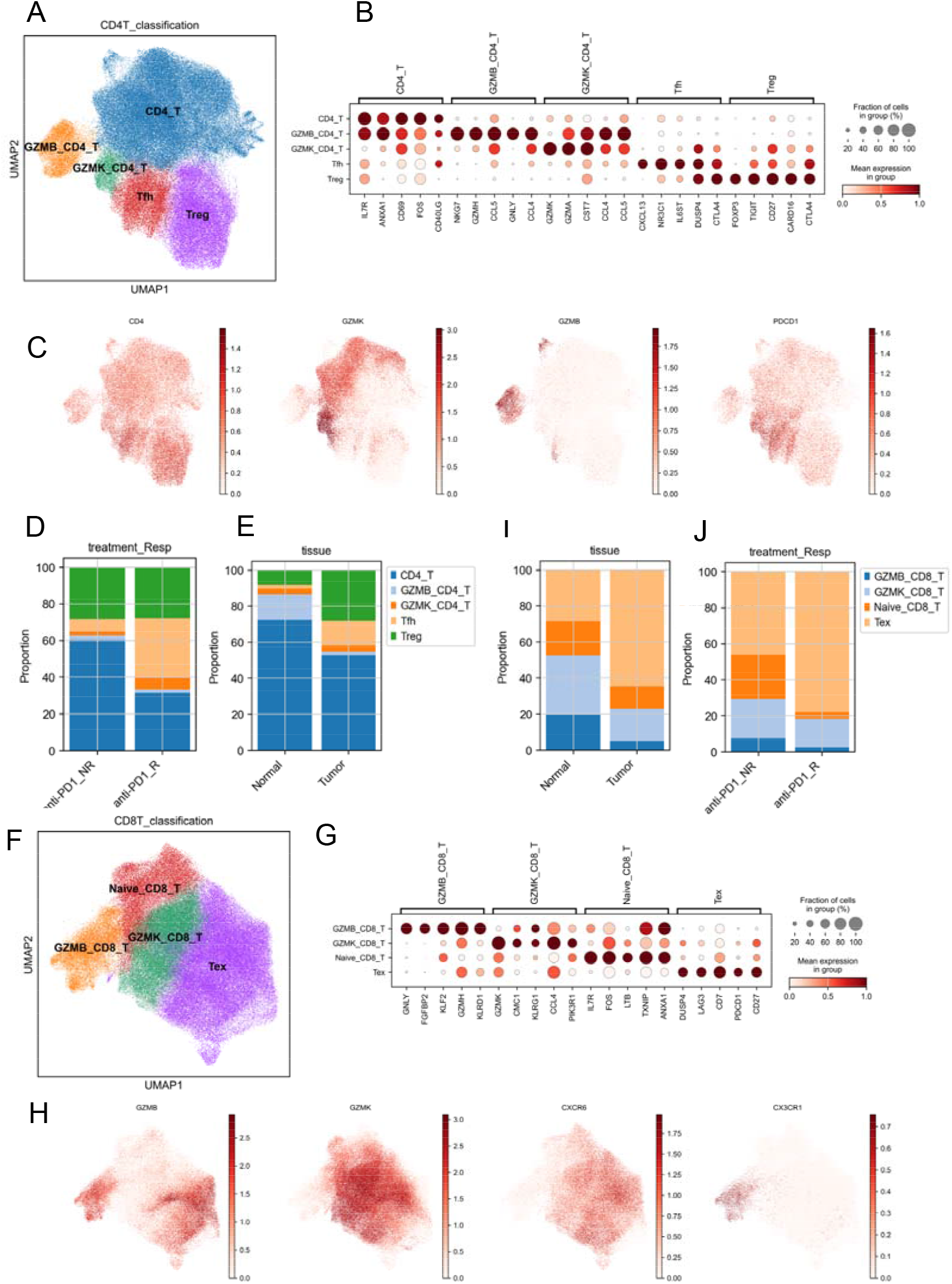
Single-cell profiling of CD4□ and CD8□ T-cell states across tissue and treatment contexts. (A) UMAP embedding of CD4□ T cells annotated by subtype. (B) Dot plot showing the average expression (color scale) and fraction of expressing cells (dot size) for selected marker genes across CD4□ T-cell subtypes in (A). (C) Feature plots displaying the expression of CD4, GZMK, GZMB, and PDCD1 overlaid on the UMAP embedding. (D) Stacked bar chart illustrating the proportions of CD4□ T-cell subtypes in anti-PD-1 non-responders (NR) versus responders (R). (E) Stacked bar chart showing the proportions of CD4□ T-cell subtypes in Normal versus Tumor samples. (F) UMAP embedding of CD8□ T cells annotated by subtype. (G) Dot plot showing the average expression (color scale) and fraction of expressing cells (dot size) for selected marker genes across CD8□T-cell subtypes in (F). (H) Feature plots displaying the expression of GZMB, GZMK, CXCR6, and CXCR1 overlaid on the UMAP embedding. (I) Stacked bar chart showing the proportions of CD8□ T-cell subtypes in Normal versus Tumor samples. (J) Stacked bar chart illustrating the proportions of CD8□ T-cell subtypes in anti-PD-1 non-responders (NR) versus responders (R).

CD8□ T cells exhibited two major differentiation trajectories (Fig. 2F-H, Supplementary Fig. 7). The first (Pathway B) progressed from naïve to GZMB□ cytotoxic T cells characterized by KLF2 and KLF3 activation, while the second (Pathway K) transitioned from naïve to GZMK□ and ultimately to exhausted (Tex) T cells via FOS/JUN-driven programs (Supplementary Fig. 7, 8). Trajectory-specific chemokine expression revealed CXCR3, CXCR6, and CCR5 enrichment along the K pathway, while CX3CR1 marked the B trajectory (Fig. 2H, Supplementary Fig. 7D). These two trajectories were balanced in normal liver but skewed toward Pathway K in tumors, implying that metabolic or hypoxic stress promotes exhaustion-prone differentiation (Fig. 2I). Consistent with this, metabolic pathway analysis showed enhanced glycolysis in exhausted T cells, whereas GZMB□ T cells relied on nitrogen metabolism and arginine biosynthesis, suggesting distinct metabolic niches underpinning their divergent fates (Supplementary Fig. 8D).

Interestingly, exhausted CD8 T cells were more abundant in responders (Fig. 2J). Given that the Magen dataset specifically focuses on PD-1 monotherapy, this finding repeatedly suggests that Tex cells represent the effector population driving the response to PD-1 immunotherapy.

### Myeloid Cells: Hidden Orchestrators of Immunotherapy Response

We characterized the myeloid compartment in liver tissues, identifying discrete subpopulations including monocytes, macrophages, Kupffer cells, and dendritic cells (DCs) (Fig. 3A-C, Supplementary Fig. 9A). Monocytes were further divided into CD14□ and CD16□ subsets, while macrophages occupied a transitional continuum bridging monocytes, Kupffer cells, and DCs. Based on transcriptional profiles, normal liver macrophages were broadly categorized into an inflammatory subset (C3, Inf_Mono_Mph_Propria), marked by IL1B expression, and a lipid-metabolism subset (C2, Lipo_Mph_Propria), expressing APOC1, TREM2, and GPNMB (Fig. 3D). The lipid-metabolism macrophages shared markers with Kupffer cells (e.g., SEPP1, EGR1, FOLR2, CD5L), suggesting they may represent a precursor state (Supplementary Fig. 9B). Kupffer cells exhibited canonical markers such as CD5L, C1QB, FTL, C1QA, and MARCO, consistent with tissue-resident specialization (Supplementary Fig. 9C)^12^.

**Fig. 3:**
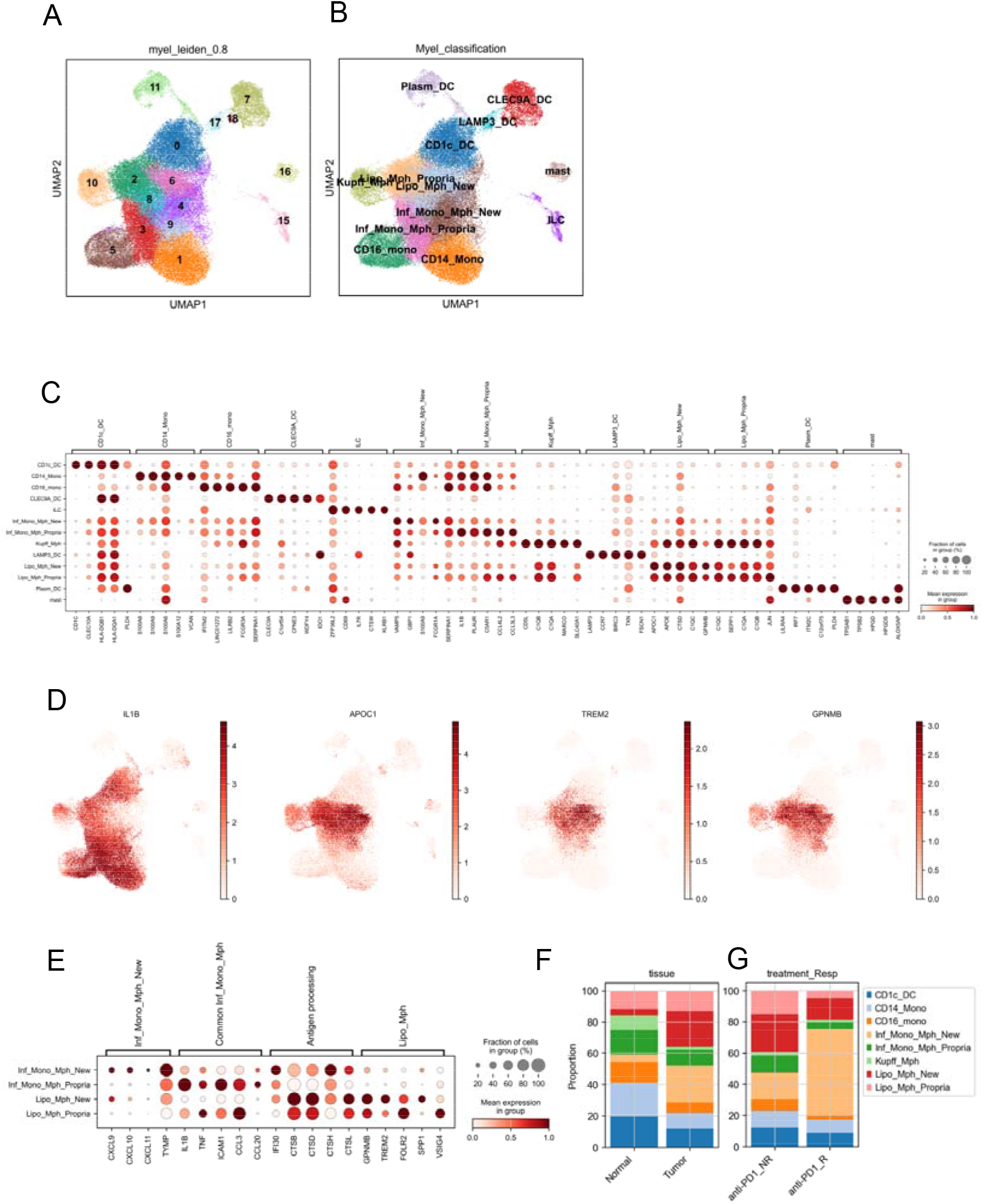
Single-cell RNA-seq profiling of myeloid cells reveals subtype diversity, marker expression patterns, and compositional shifts across tissue states and anti-PD-1 treatment response. (A) UMAP embedding of myeloid cells after scVI integration for batch correction, clustered using the Leiden algorithm (resolution = 0.8). (B) UMAP visualization of the same cells annotated with refined myeloid cell identities. (C) Dot plot showing the average expression (color scale) and fraction of expressing cells (dot size) for marker genes across annotated myeloid subtypes. (D) Feature plots displaying the expression of IL1B, APOC1, TREM2, and GPNMB overlaid on the UMAP embedding. (E) Dot plot displaying the average expression (color scale) and fraction of expressing cells (dot size) for genes representing functional modules across macrophage subtypes. (F) Stacked bar chart illustrating the relative proportions of myeloid subtypes in Normal versus Tumor samples. (G) Stacked bar chart showing the distribution of myeloid subtypes in anti-PD-1 non-responders (NR) and responders (R).

Dendritic cells were subdivided into cDC1 (CLEC9A □), cDC2 (CD1C □), plasmacytoid DCs (pDCs, LILRA4 □, IRF7 □), and LAMP3 □ DCs, a recently described population expressing CCR7, LAMP3, CCL19, CXCL9, CD80, and CD86, potentially involved in lymphoid follicle formation and antigen presentation (Fig. 3C)^36^. Other innate populations included ILCs (KIT □, AREG □, GATA3 □) and mast cells (KIT □, MS4A2 □) (Supplementary Fig. 9D).

Comparative analysis of tumor versus normal tissue revealed a significant reduction of Kupffer cells within the tumor microenvironment, accompanied by dynamic changes in macrophage subpopulations. Tumors exhibited expansion of two distinct populations: a novel inflammatory macrophage subset (Inf_Mono_Mph_New, IMN), expressing CXCL9/10/11, IL1B, CCL3, and CCL20, and a new lipid-metabolism subset (Lipo_Mph_New, LMN), marked by GPNMB and SPP1. IMN cells displayed both antigen-processing capacity (IFI30, CTSB) and T-cell chemoattractant activity, whereas LMN cells retained antigen-processing function but lacked T-cell recruitment features, suggesting a potential role in immune tolerance (Fig. 3E).

Stratifying tumors by response to immunotherapy revealed that responders exhibited pronounced enrichment of the IMN population (Fig. 3G). This suggests that IMN macrophages may serve as critical modulators of PD-1 therapy efficacy, representing a “hidden driver” of antitumor immune activity. Collectively, these findings highlight the heterogeneity and functional specialization of myeloid cells in the liver tumor microenvironment and underscore their potential impact on immunotherapeutic outcomes.

### Single-Cell Profiling Reveals B Cell Heterogeneity and Functional States in the Liver Tumor Microenvironment

We next focused on B cells using the Magen dataset. B cells in the liver microenvironment can be broadly categorized into three major groups: naïve B cells, memory B cells, and plasma cells (Fig. 4A, B). Within the naïve compartment (IGHD □), two distinct subpopulations were identified: resting naïve B cells (C4) and early-activated B cells (CD69 □ CD83 □; C0, 5, 11). The early-activated subset retains IGHD expression while upregulating activation markers CD69 and CD83, indicating that IGHD silencing occurs after activation (Fig. 4C). Notably, a CCR7 □ subset, which is more abundant in normal liver, downregulates CCR6 and CXCR5 while maintaining CXCR4, suggesting migration from the T-cell zone toward CXCL12-rich regions such as the Kupffer cell area (Fig. 4D, Supplementary Fig. 10A). Given the co-expression of CD69 and CD83 but lack of CD80/CD86, these cells likely represent plasmablast-like precursors contributing to hepatic immune tolerance. This is consistent with CXCL12–CXCR4 signaling supported by Kupffer cells and hepatic stellate cells, which may guide activated B cells into the hepatic parenchyma (Supplementary Fig. 10B, C).

**Fig 4.**
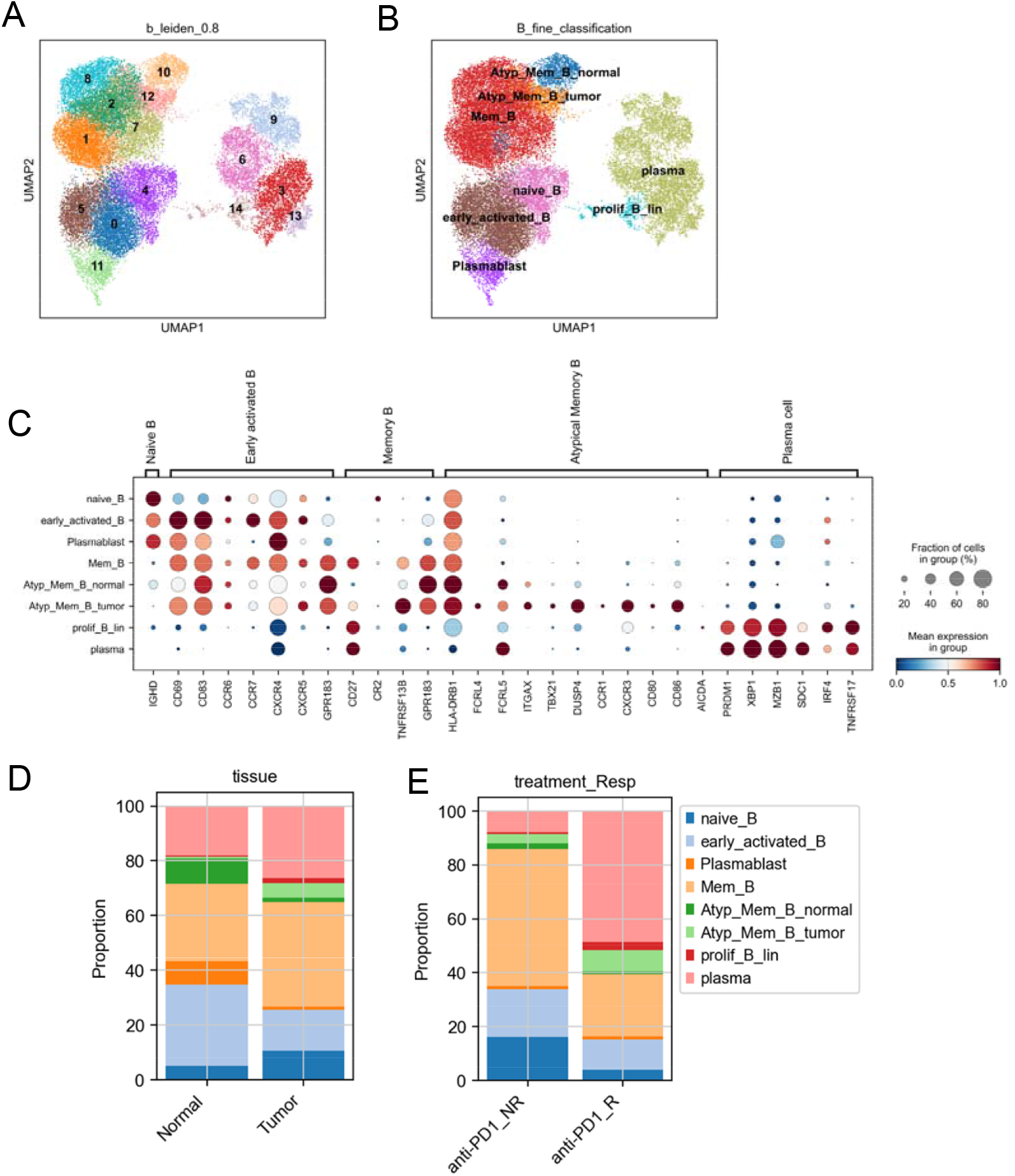
Characterization of B-cell heterogeneity in the tumor microenvironment and its association with anti-PD-1 therapy response. (A) UMAP embedding of B cells from the scVI-integrated dataset (batch-corrected), clustered using the Leiden algorithm (resolution = 0.8). (B) UMAP visualization of B cells annotated with refined B-cell subtype identities. (C) Dot plot displaying the average expression (color) and fraction of expressing cells (dot size) for selected marker genes across B-cell subtypes. (D) Stacked bar chart illustrating the relative proportions of B-cell subtypes in Normal and Tumor tissues. (E) Stacked bar chart showing the distribution of B-cell subtypes in anti-PD-1 therapy non-responders (NR) and responders (R).

Memory B cells (IGHD □, CD27 □, TNFRSF13B □, CD86 □) were further divided into classical and atypical (or follicular-peripheral) subtypes. The atypical memory B cells (AMB; C10/C12) are characterized by downregulation of CD27 and CD21, and upregulation of HLA-DRB1, ITGAX, TBX21, DUSP4, and FCRL5. Notably, tumor-associated AMB (C12) lose IGHD expression and gain FCRL4, along with increased CXCR3, CCR1, and CXCR5 levels, while reducing CCR7 (Supplementary Fig. 10A). These cells exhibit elevated CD80 and CD86, suggesting enhanced T-cell costimulatory potential. Since responder samples contained more C12 cells, this subset may play a pivotal role in antitumor immunity through T-cell activation (Fig. 4E). Co-expression of AICDA in C12 and proliferating clusters implies that these cells engage in somatic hypermutation and class switching, analogous to germinal center light-zone B cells undergoing affinity maturation under T-cell help.

Another tumor-enriched population (C7) upregulates cytoskeletal regulators (ACTB, ACTG1, COTL1, RAC2) while downregulating key homing receptors (CCR7, CXCR4, CXCR5, GPR183), indicating a highly migratory phenotype. The overlap of migration-associated markers between C7 and C12 suggests that both subsets acquire motility after completing T-cell interactions, potentially contributing to the spatial remodeling of the B-cell compartment in tumors. A separate cluster (C8), enriched in tumor upregulates stress-response genes (DNAJB1, HSPA1A, HSP90AA1, HSPE1), reflecting the metabolic or inflammatory stress within the tumor niche (Supplementary Fig. 10A, C).

Finally, plasma cells are defined by high expression of SDC1, PRDM1, XBP1, and MZB1 (Fig. 4A). These cells are enriched in responders and upregulate CXCR3, linking them to CXCL9/10/11-producing macrophages that may mediate long-term retention within the hepatic tumor microenvironment (Fig. 3E, 4C, Supplementary Fig. 10F). An intermediate subset (C6) expresses PRDM1 and IRF4 but shows lower TNFRSF17 (BCMA), representing a transitional stage in plasma cell differentiation (Supplementary Fig. 10F). Together, these findings reveal a continuum of B-cell states in the liver, spanning from naïve activation and migration to memory formation and terminal plasma differentiation, intricately shaped by spatial cues and local immune interactions.

### Chemokine-Receptor Interactions and Immune Regulation in the Tumor Microenvironment

By analyzing chemokine-receptor interactions, we aimed to uncover the immune recruitment dynamics distinguishing tumor from normal liver tissue. Myeloid cells emerge as pivotal orchestrators of immune cell recruitment within the tumor microenvironment. The IMN subset, characterized by robust secretion of chemokines CXCL9, CXCL10, and CXCL11, likely promotes the migration of CXCR3-expressing lymphocytes, including T cells, memory B cells, and plasma cells (Fig. 3E, 5A, supplementary Fig. 10F). Notably, T regulatory cells (Tregs) also upregulate CXCR3, suggesting a competitive chemotactic interplay that may contribute to immune tolerance. In parallel, both IMN and LAMP3□ dendritic cells upregulate CCL19, the ligand for CCR7 (Fig. 5B). CCR7 is expressed on naïve T cells as well as B cells, suggesting that this chemokine-receptor axis may recruit lymphocytes and contribute to the formation of tertiary lymphoid structures within the liver and tumor microenvironment. Additionally, the LMN myeloid subset secretes CCL8 and CCL18, chemokines capable of recruiting CCR8□ Tregs, suggesting a myeloid-mediated mechanism for local immunosuppression and modulation of antitumor immunity (Fig. 5C).

**Fig. 5:**
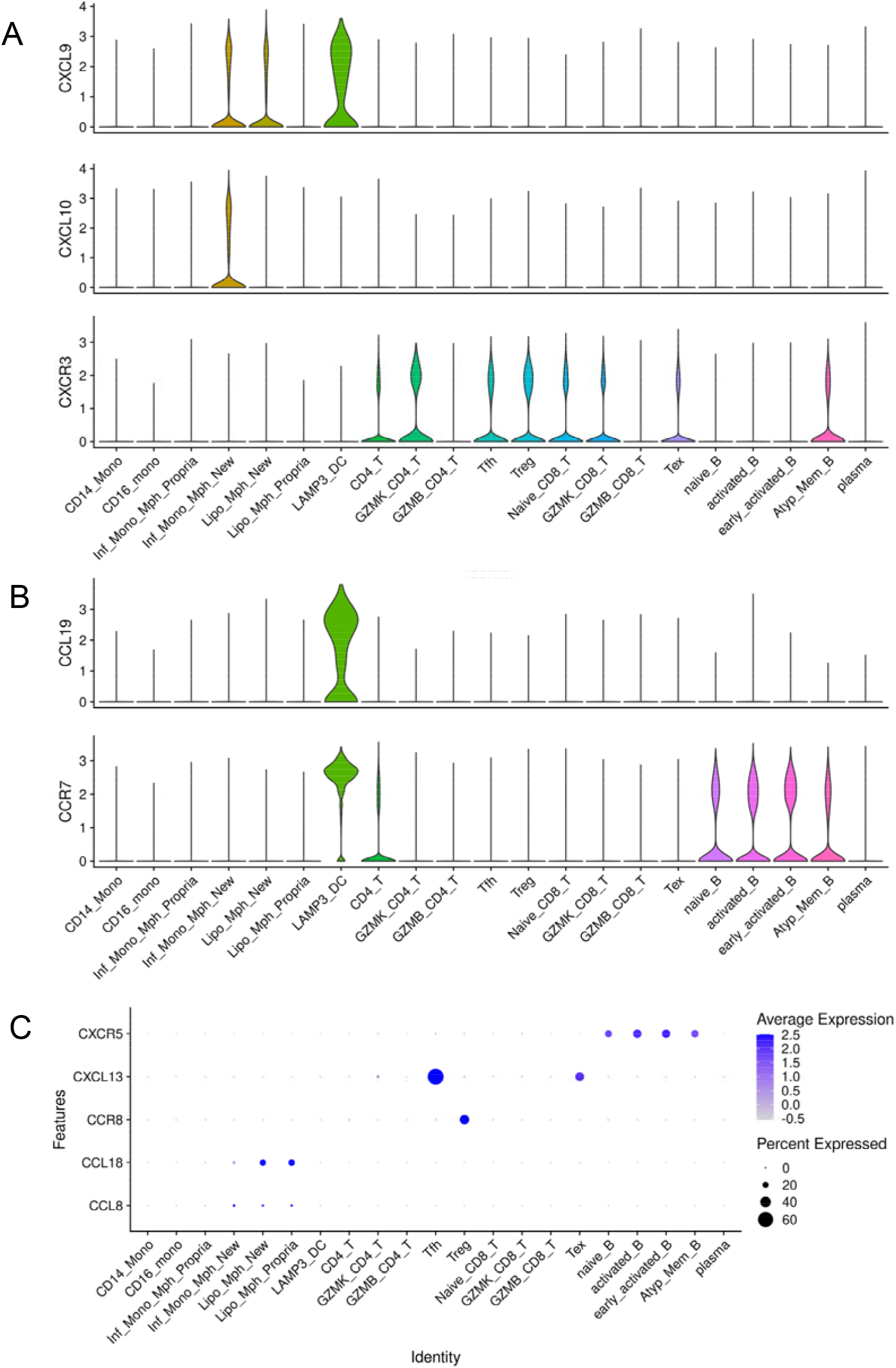
Expression patterns of selected chemokine ligands and receptors across immune cell populations. (A) Violin plots showing the expression of the CXCL9–CXCL10–CXCR3 ligand–receptor axis across immune cell populations; violin width reflects the density of cells at each expression level. (B) Violin plots showing the expression of the CCL19–CCR7 ligand–receptor pair across immune cell populations. (C) Dot plot displaying the average expression (color scale) and fraction of expressing cells (dot size) for five genes (CXCR5, CXCL13, CCR8, CCL18, and CCL8) across immune cell populations.

**Figure 6.**
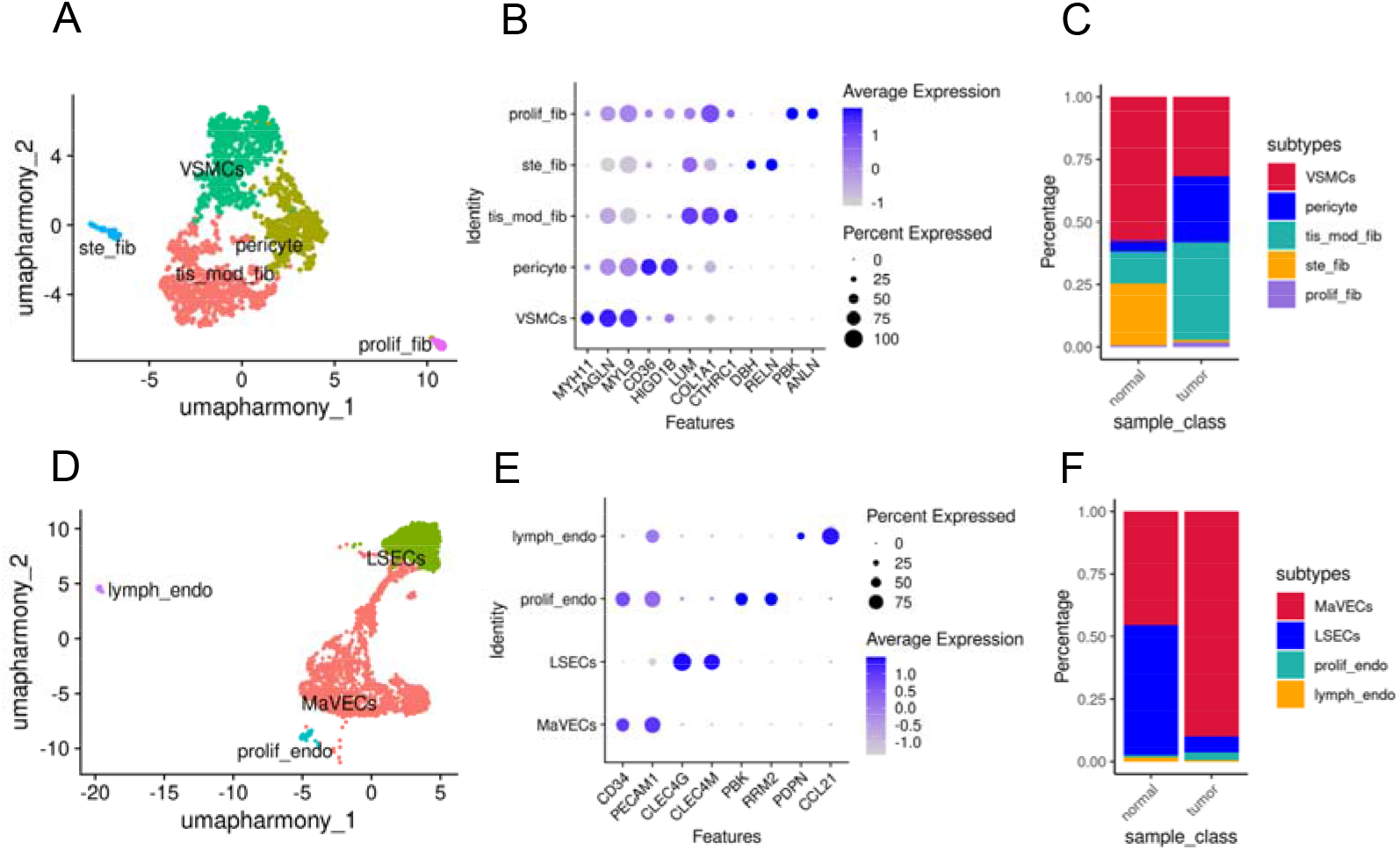
Single-cell characterization of stromal cell subtypes in normal and tumor tissues. (A) UMAP visualization of fibroblasts (combinedfrom tumor and normal samples) annotated by cell subtype identity. (B) Dot plot showing the average expression (color scale) and fraction of expressing cells (dot size) for selected marker genes across fibroblast subtypes in (A). (C) Stacked bar chart illustrating the relative proportions of fibroblast subtypes in Normal versus Tumor samples. (D) UMAP embedding of endothelial cell populations (combined from normal and tumor samples) annotated by cell subtype identity. (E) Dot plot showing the average expression (color scale) and fraction of expressing cells (dot size) for selected marker genes across endothelial subtypes in (D). (F) Stacked bar chart illustrating the relative proportions of endothelial subtypes in Normal versus Tumor samples.

Additionally, we found evidence that T follicular helper (Tfh) cells actively participate in the formation of lymphoid follicles. Rather than following the canonical paradigm in which Tfh upregulate CXCR5 and migrate toward CXCL13-rich regions, our data reveal an alternative pattern: Tfh cells themselves exhibit reduced CXCR5 expression but increased CXCL13 production (Fig. 5C)^37^. This suggests that Tfh cells may actively recruit CXCR5□ B cells to their vicinity, thereby driving the local assembly of follicular structures in situ.

Together, these observations highlight how immune cells coordinate chemokine-receptor interactions to sculpt the immune landscape of tumors, balancing lymphocyte recruitment, activation, and local tolerance.

### Stromal Compartment: Fibroblasts and Endothelial Cells

Stromal cells (CD45 □) were analyzed using the Lu dataset^4^. To compare the stromal microenvironment between normal and tumor tissues, fibroblasts derived from both conditions were jointly clustered. Seurat analysis identified five major stromal populations, annotated as hepatic stellate cells (ste_fib), vascular smooth muscle cells (VSMCs), pericytes, tissue-remodeling fibroblasts (tis_mod_fib), and proliferative fibroblasts (Fig. 5A, B). Notably, hepatic stellate cells were significantly reduced in tumor tissues, whereas pericytes and tissue-remodeling fibroblasts were markedly enriched (Fig. 5C). Proliferative fibroblasts were rare in normal liver but expanded in tumors, indicating an active stromal proliferative state within the tumor microenvironment.

We next examined endothelial cell populations in tumor tissues. Consistent with the hypervascular nature of hepatocellular carcinoma, liver sinusoidal endothelial cells (LSECs) were markedly decreased, while macro-vascular endothelial cells (MaVECs) were significantly increased, reflecting a shift toward a vascular architecture characteristic of tumor-associated neovascularization and enhanced blood supply^11^.

## Discussion

Hepatocellular carcinoma (HCC) is a common disease worldwide, and numerous genomic and transcriptomic studies have investigated this malignancy ^4,9,24^. However, systematic descriptions of transcriptional programs in HCC remain limited. Here, using established algorithms (e.g., Harmony and scVI), we repeatedly identified consistent transcriptional patterns in malignant cells, T cells, and myeloid cells across multiple datasets. Although our cell-level annotations may not be highly granular, these patterns are reproducible across independent cohorts.

Encouragingly, we identified reproducible transcriptional programs in malignant hepatocytes. At the transcriptomic level, tumor cell dedifferentiation appears to follow a “three-step” trajectory. Notably, malignant progenitor–like cells are already present at early stages. Although cancers display substantial heterogeneity at the genomic level, they appear to converge on shared features at the RNA level, a phenomenon also reported in other tumor types ^38^. Importantly, genes such as TFF2 and TFF3, which represent gastrointestinal lineage signals, play a prominent role during hepatocellular carcinoma dedifferentiation (Supplementary Data 1). These dedifferentiation processes and RNA-level alterations may provide potential opportunities for RNA-based therapeutics.

We identified a population of GZMB □ CD4 T cells enriched in normal liver tissue. These cells likely represent cytotoxic CD4 T cells, a tissue-resident immune subset involved in local immune surveillance. The liver is constantly exposed to gut-derived antigens and requires rapid but tightly controlled immune responses. Cytotoxic CD4 T cells may provide a low-inflammatory mechanism for eliminating infected or damaged hepatocytes. Their reduction in tumor tissue suggests that hepatocellular carcinoma disrupts this resident immune surveillance niche^39^.

CD8 T-cell differentiation appears to be bidirectional. In the dataset from Liu et al., this population was termed FGFBP2 □ NK-like T cells^10^; however, these cells clearly express TCR and are therefore bona fide T cells. Previous studies have classified CD8 T cells into GZMK □ and GZMB □ subsets based on granzyme expression^40^. Based on our single-cell analyses, we propose that CD8 T cells follow two differentiation trajectories. One trajectory gives rise to GZMK □ GZMB □ cells. The other originates from naïve T cells that differentiate into GZMK □ cells, which subsequently progress toward exhaustion and gradually upregulate GZMB as the exhausted phenotype intensifies. This finding suggests that GZMB□ cells have dual origins, indicating that classification of CD8 T cells solely by GZMK/GZMB expression is overly simplistic. Although both Tex and GZMB □ CD8 T cells express GZMB, they exhibit marked transcriptional differences. GZMB □ CD8 T cells do not express inhibitory molecules such as PDCD1 and LAG3, whereas Tex cells lack expression of GZMB-subset markers such as CX3CR1, FGFBP2, and FCGR3A. We reproduced this CD8 T-cell stratification across multiple datasets, suggesting that this mode of T-cell classification represents a broadly conserved principle.

This study has several limitations. First, our analyses are based on retrospective, publicly available datasets without an independent validation cohort, and therefore the clinical and mechanistic implications require prospective confirmation. Second, the conclusions rely primarily on single-cell RNA sequencing, which lacks spatial context; thus, inferred cell–cell interactions and immune recruitment dynamics will need validation using spatial transcriptomics or multiplex imaging. Finally, as with all dissociation-based single-cell studies, residual technical artifacts such as doublets cannot be completely excluded and may influence the interpretation of rare or transitional cell states.

## Data and Code Availability

All raw data used in this study were obtained from public databases: Lu (GSE149614), Magen (GSE206325), Sharma (GSE156337), and Liu (GSE243013). All methods were based on previously published and publicly available algorithms.

## Acknowledgements

We thank the authors for generously providing the original data. We also thank the developers who shared their algorithms online.

