## Supplementary Figure 1-4 for "Hepatocellular Carcinoma Revisited from Single Cell Sequencing: Dynamic Evolution from Epithelial Dedifferentiation to Mesenchymal Remodeling"

### Slide 1
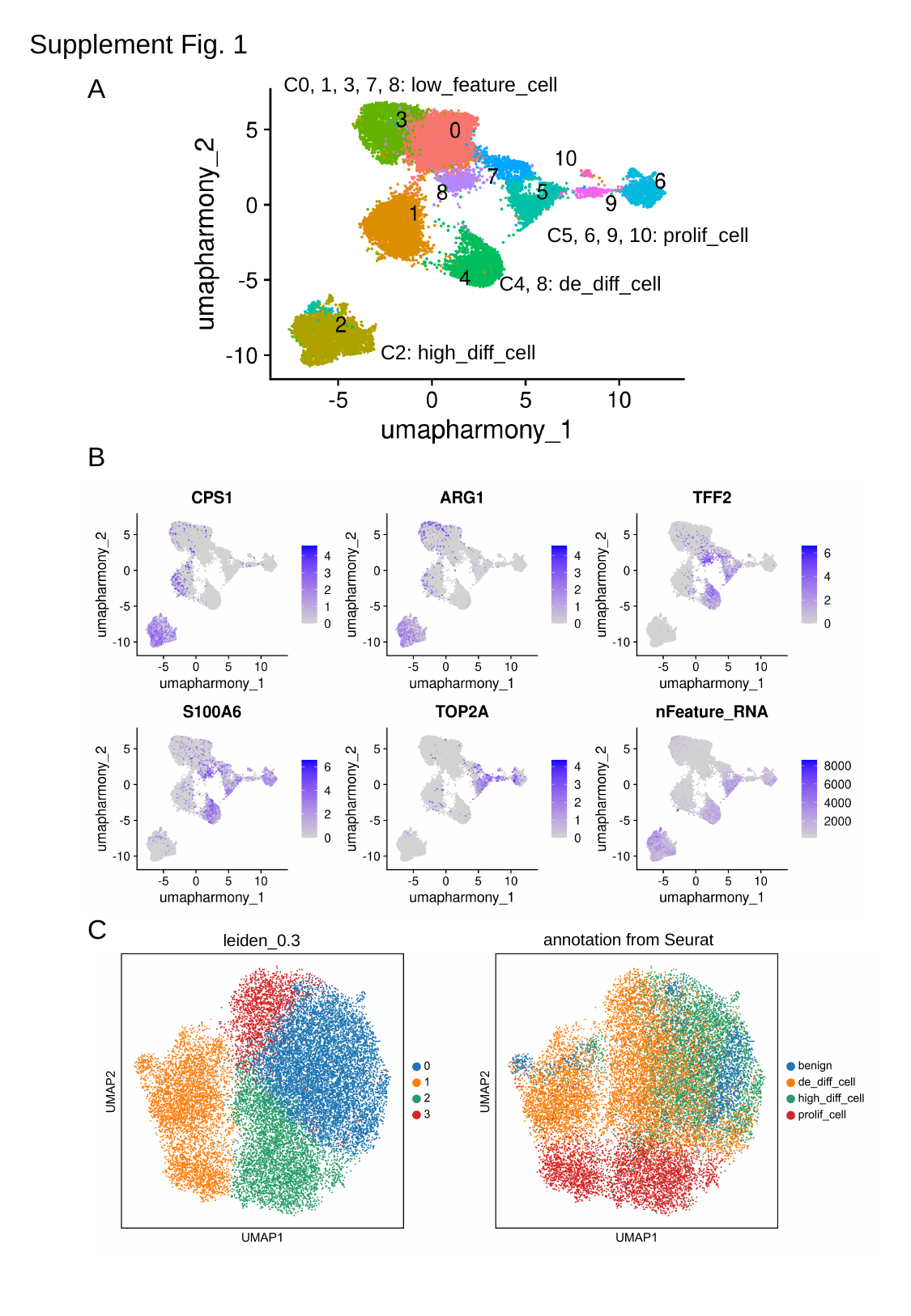

Supplement Fig. 1
A
C0, 1, 3, 7, 8: low_feature_cell
C5, 6, 9, 10: prolif_cell
C4, 8: de_diff_cell
C2: high_diff_cell
B
C
annotation from Seurat
leiden_0.3

### Slide 2
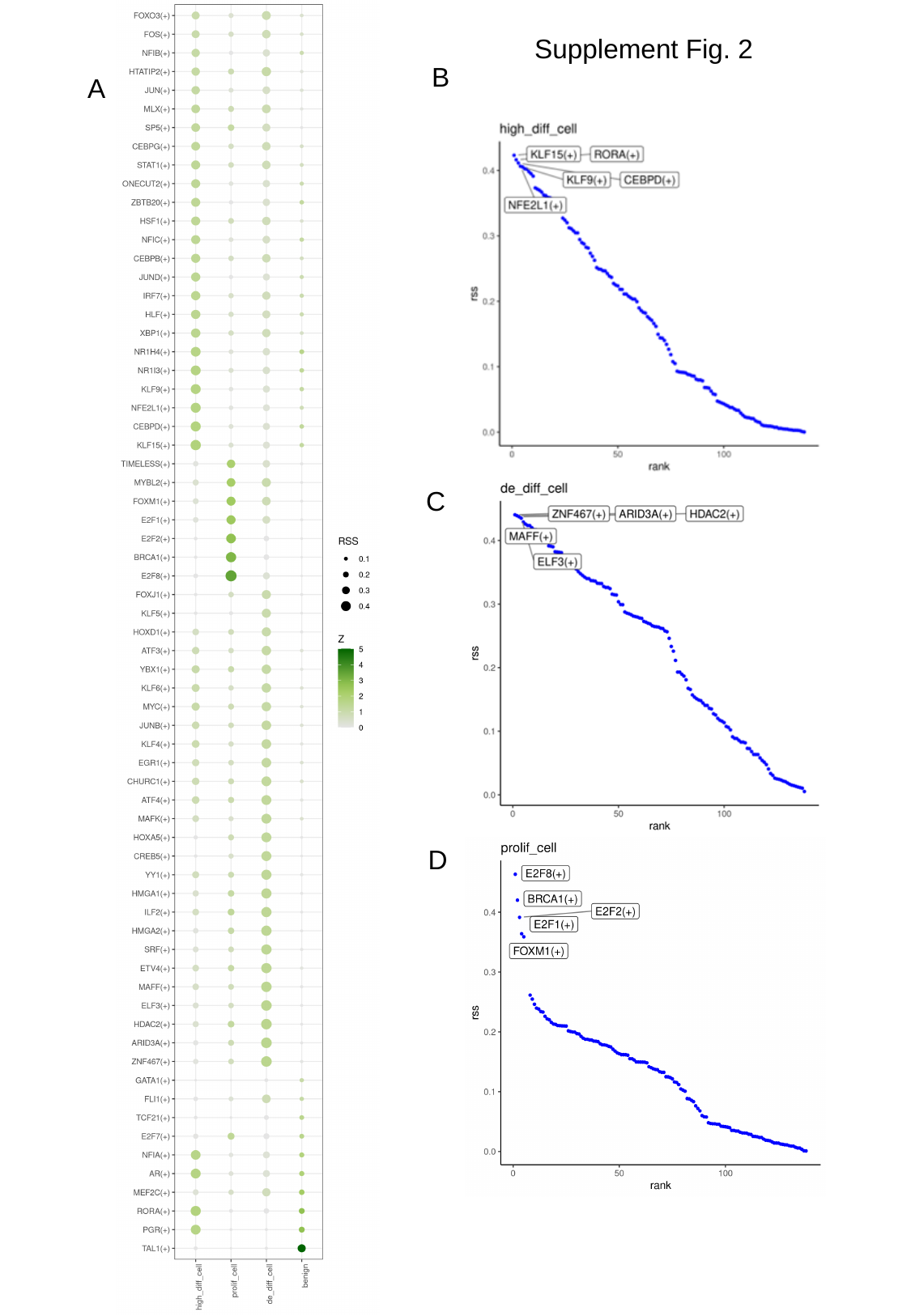

Supplement Fig. 2
B
A
C
D

### Slide 3
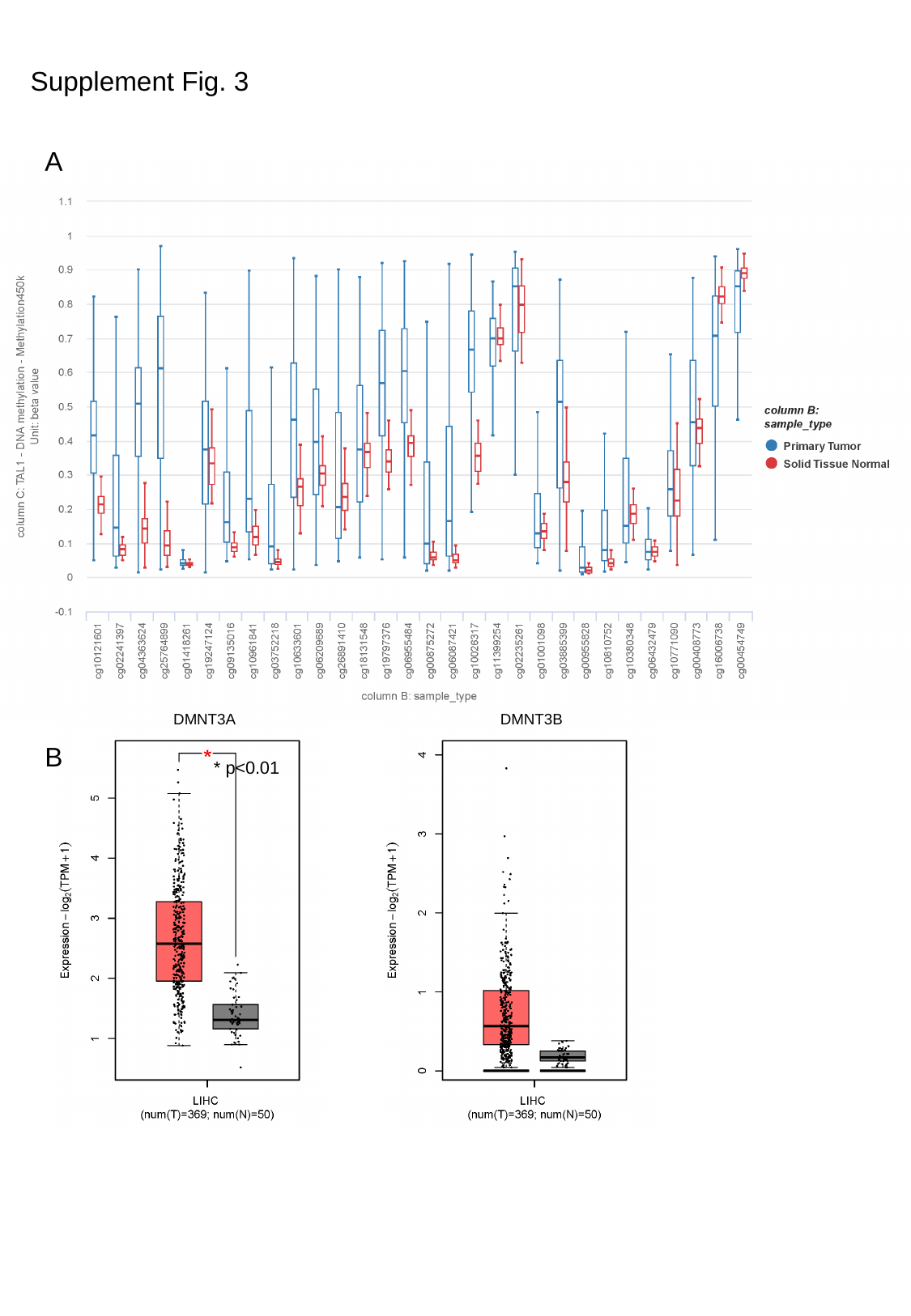

Supplement Fig. 3
A
DMNT3A
DMNT3B
B
* p<0.01

### Slide 4
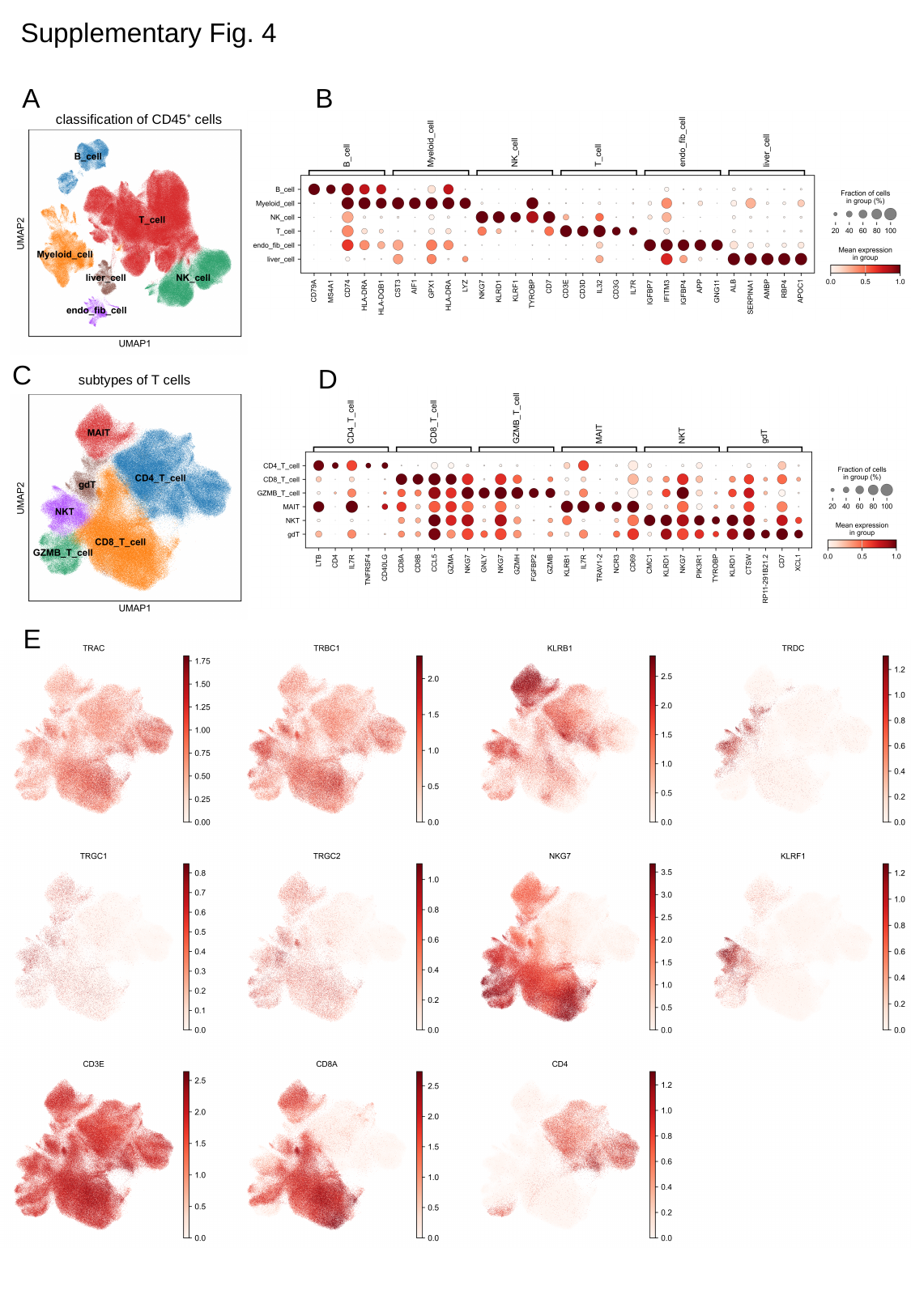

Supplementary Fig. 4
A
B
classification of CD45⁺ cells
C
D
subtypes of T cells
E
