## Supplementary Figure 5-8 for "Hepatocellular Carcinoma Revisited from Single Cell Sequencing: Dynamic Evolution from Epithelial Dedifferentiation to Mesenchymal Remodeling"

### Slide 1
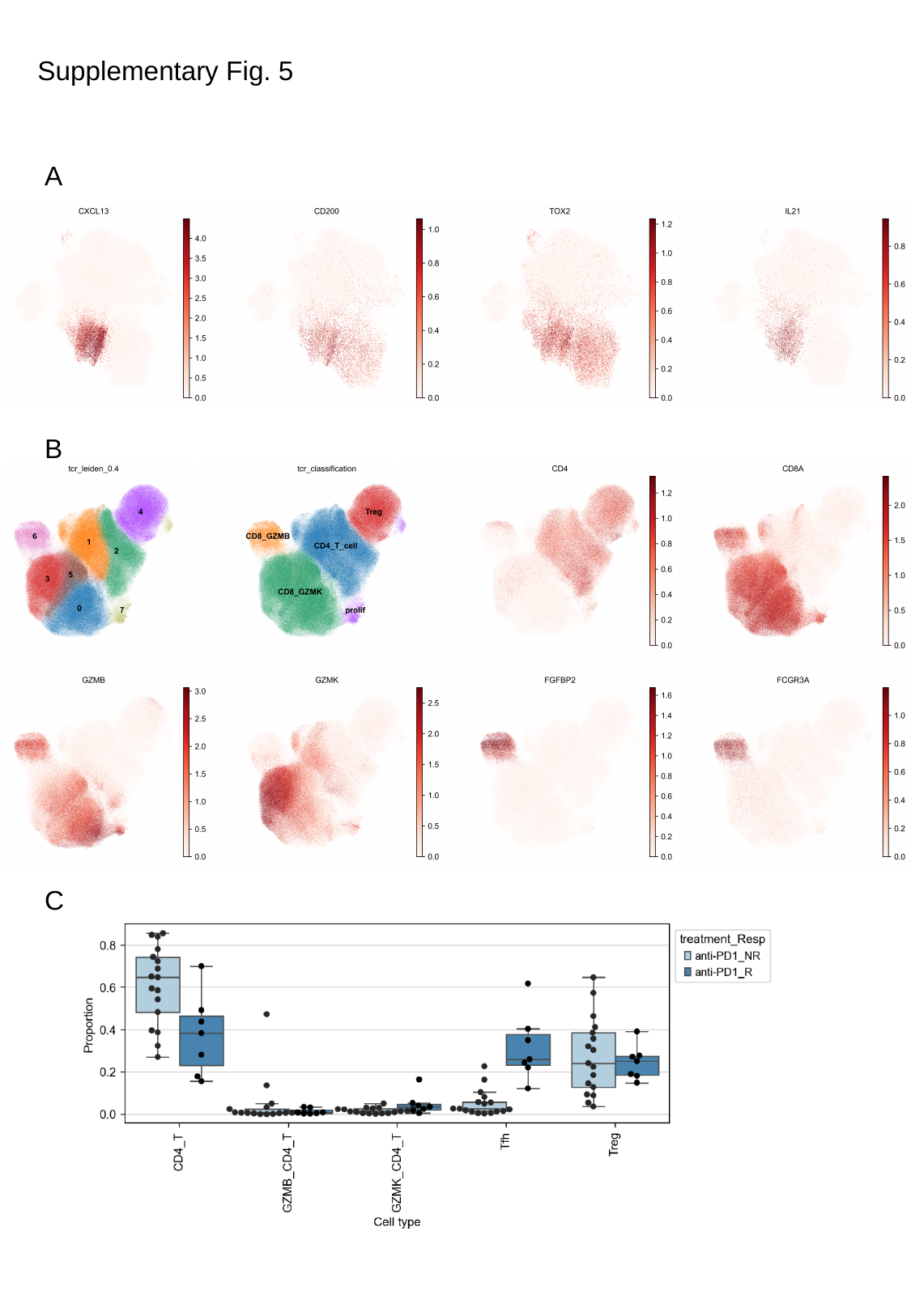

Supplementary Fig. 5
A
B
C

### Slide 2
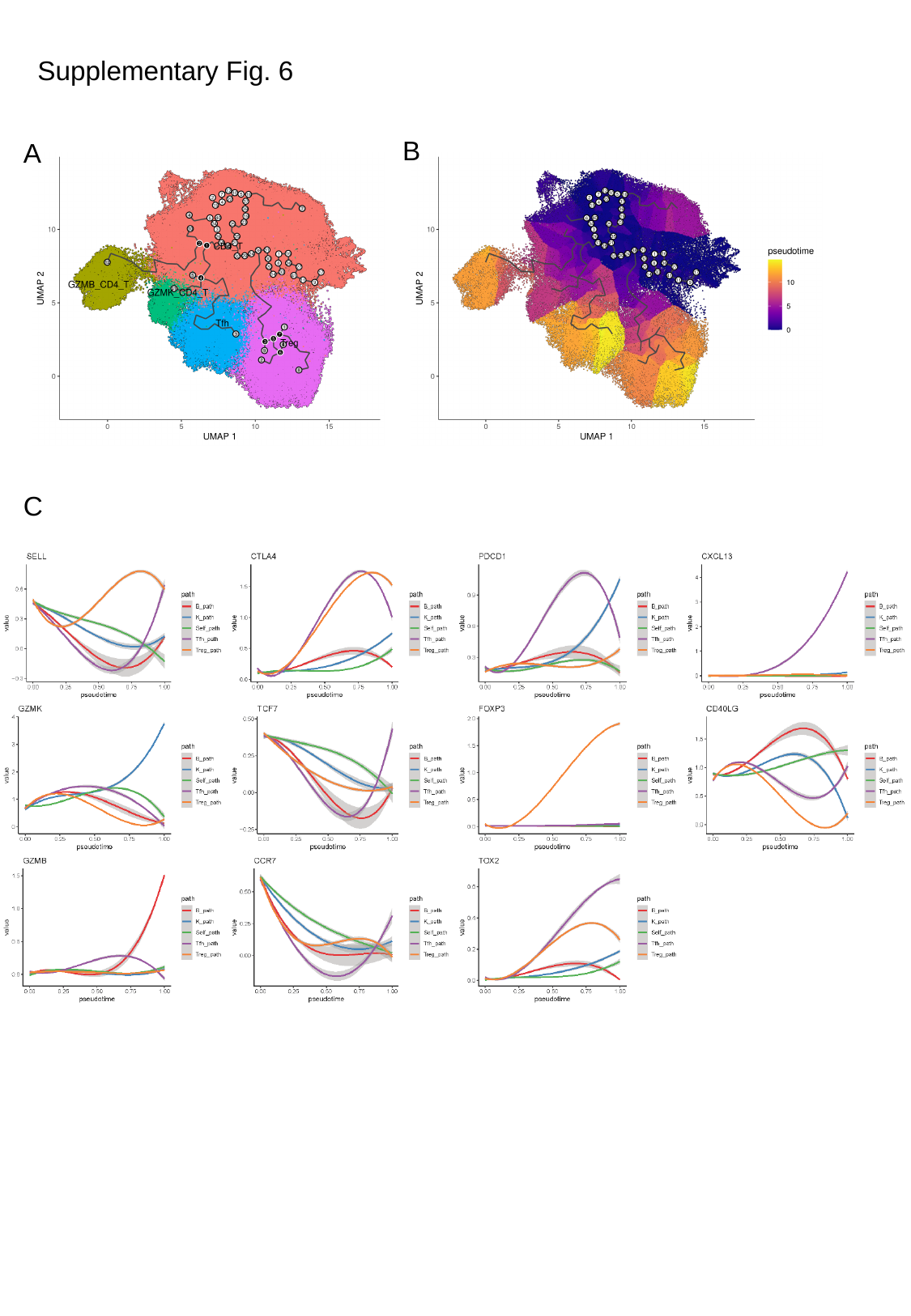

Supplementary Fig. 6
B
A
C

### Slide 3
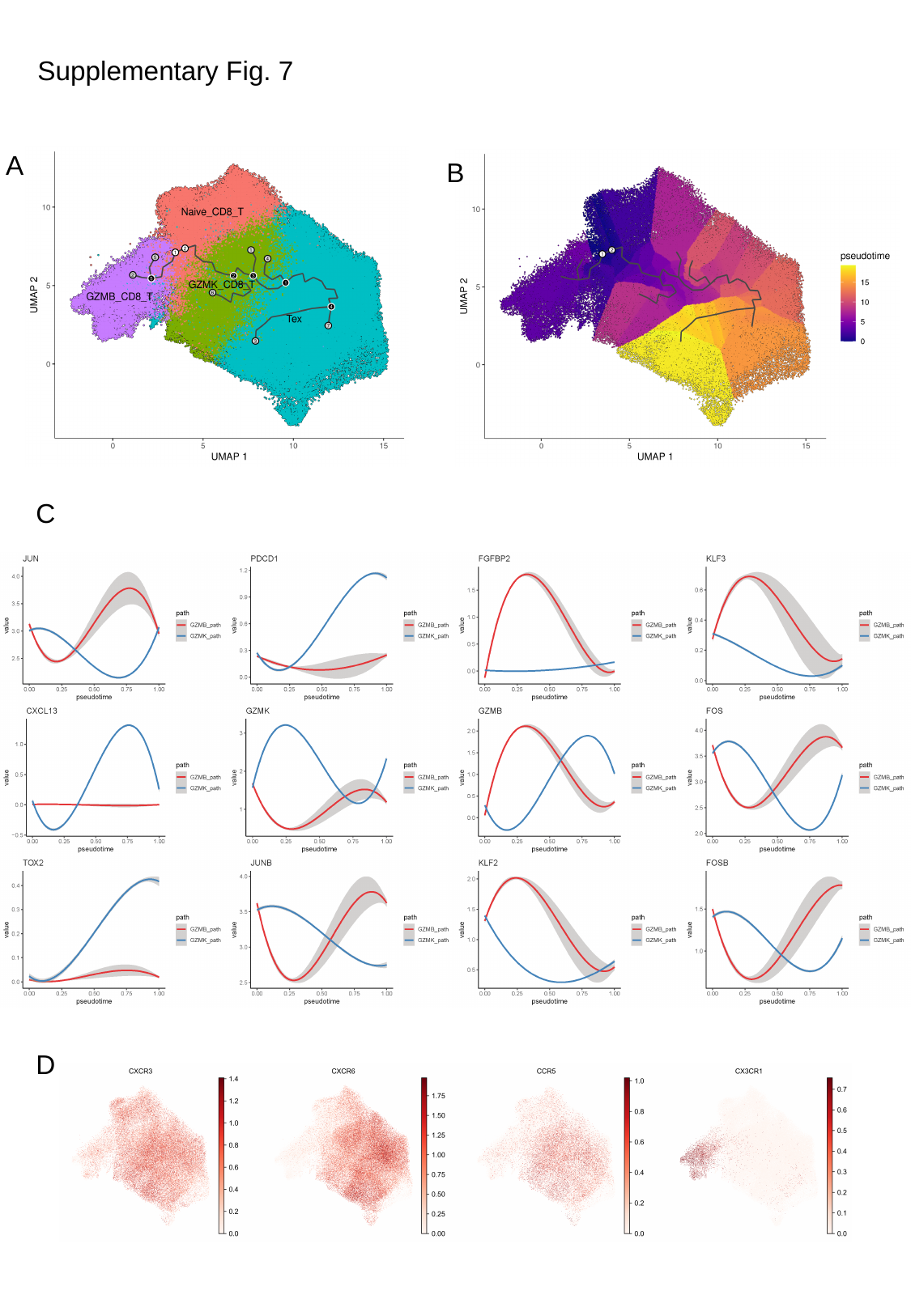

Supplementary Fig. 7
A
B
C
D

### Slide 4
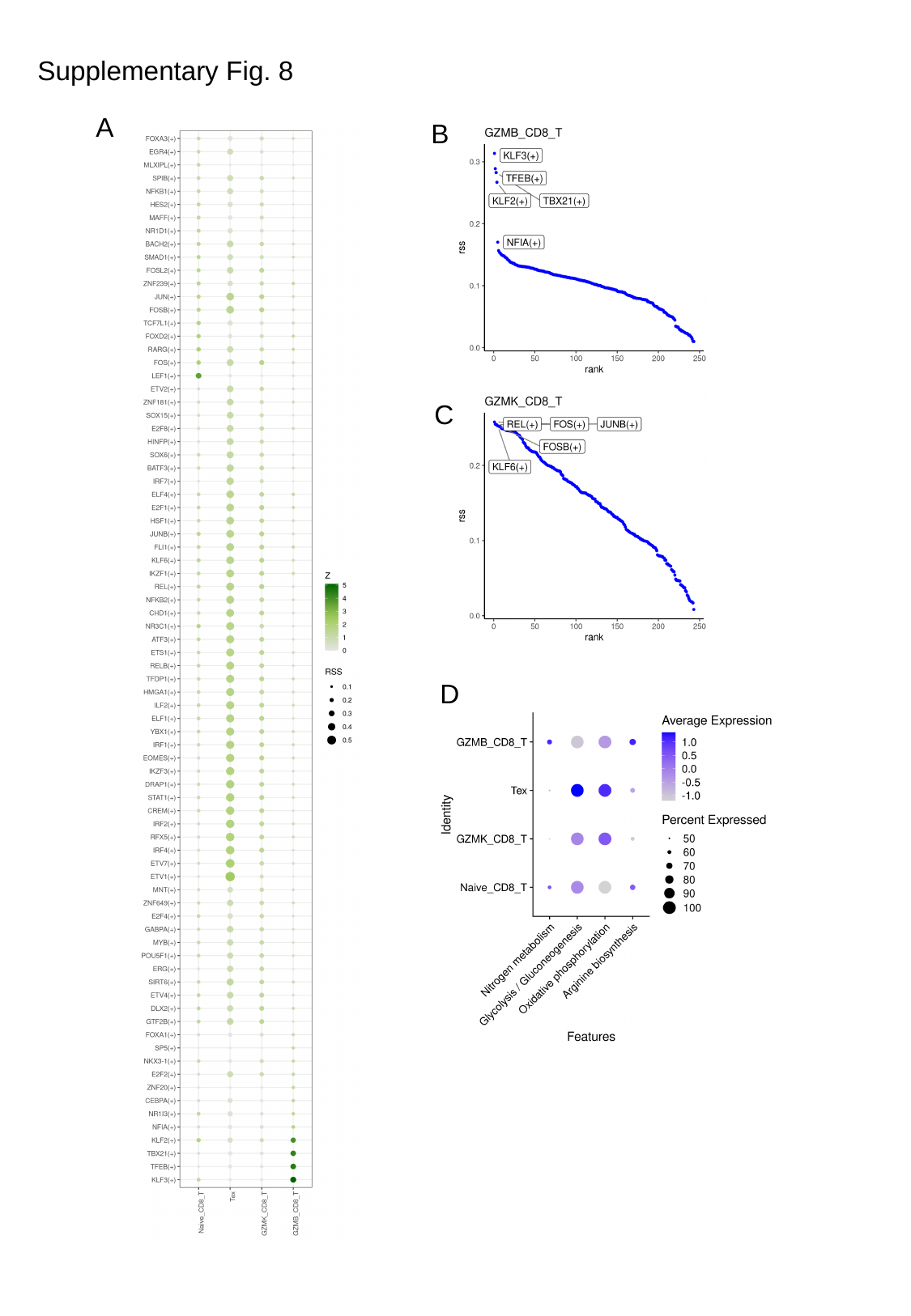

Supplementary Fig. 8
A
B
C
D
