## Supplementary figures and images for "Hepatocellular Carcinoma Revisited from Single Cell Sequencing: Dynamic Evolution from Epithelial Dedifferentiation to Mesenchymal Remodeling"

### Supplementary Figure 9-10

#### Slide 1
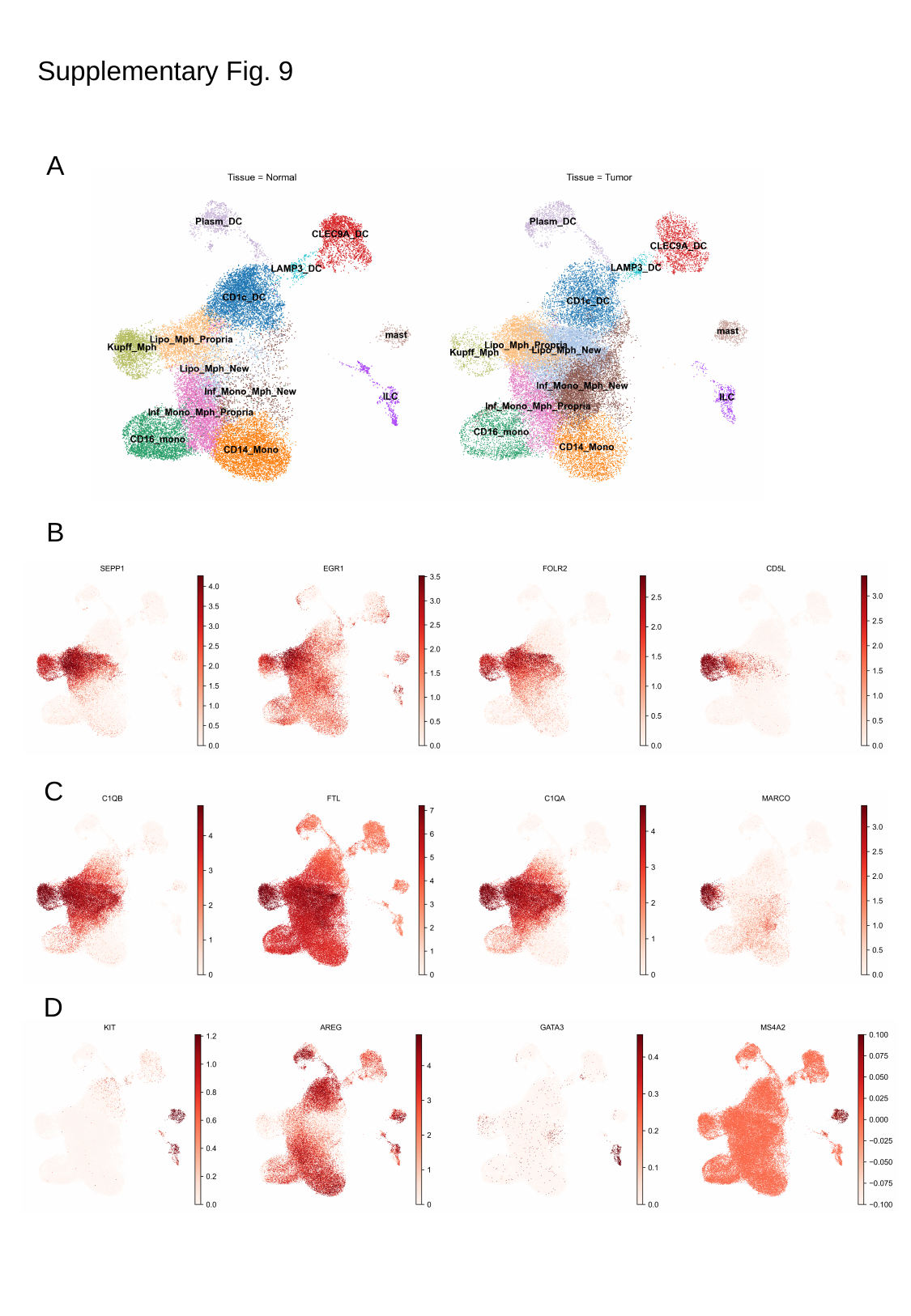

Supplementary Fig. 9
A
B
C
D

#### Slide 2
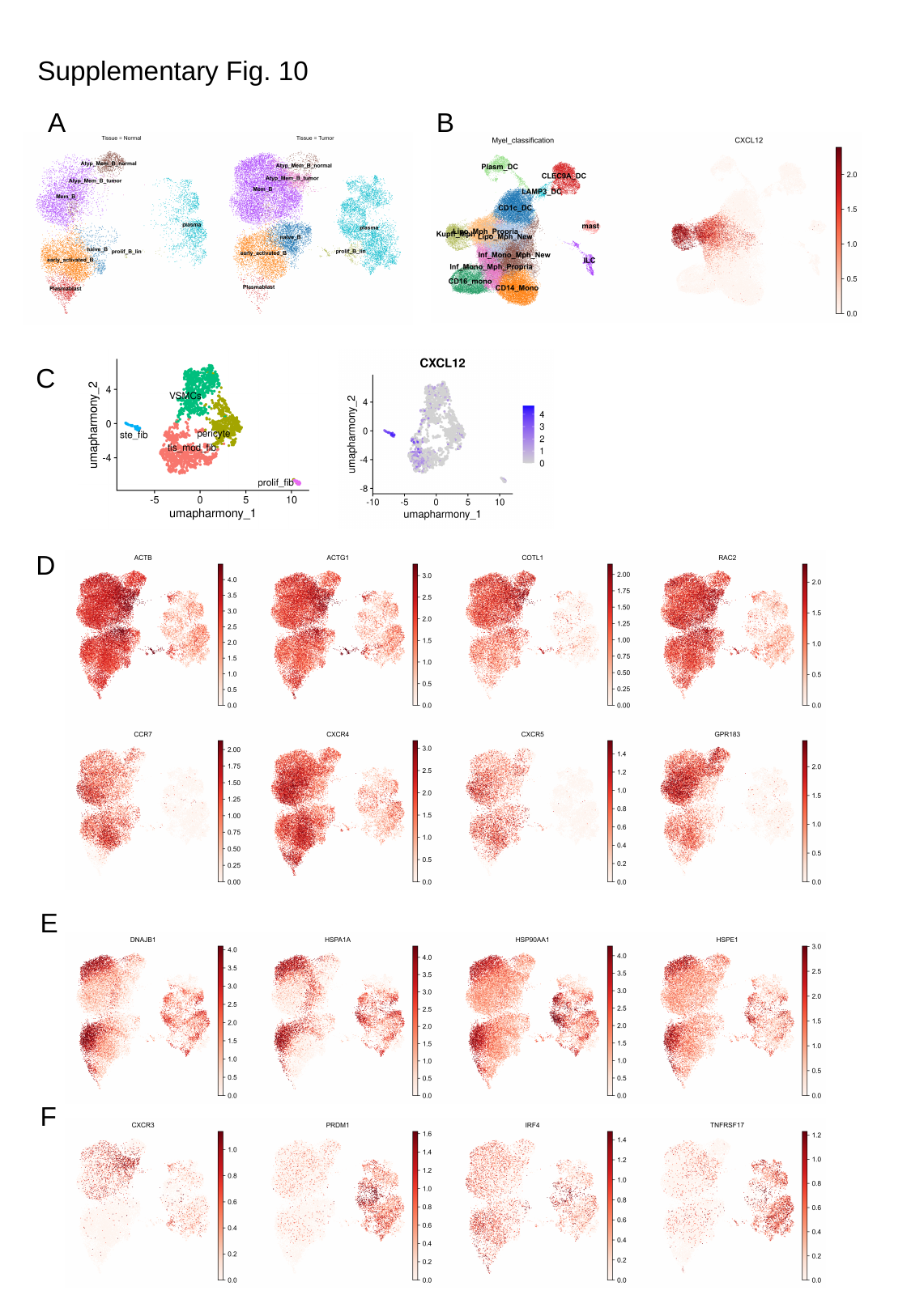

Supplementary Fig. 10
A
B
C
D
E
F
