## Supplementary Figure Legends for "Hepatocellular Carcinoma Revisited from Single Cell Sequencing: Dynamic Evolution from Epithelial Dedifferentiation to Mesenchymal Remodeling"

### Legends of Supplementary Figures

#### Supplementary Figure 1: Validation of epithelial three-step de-differentiation across datasets and methods.

(A) UMAP of integrated epithelial cells (batch effects corrected using Harmony) clustered using the Louvain algorithm (resolution = 0.5). Clusters are annotated based on functional states: C0, C1, C3, C7, and C8 as low_feature_cell; C5, C6, C9, and C10 as prolif_cell; C4 and C8 as de_diff_cell; and C2 as high_diff_cell.

(B) Feature expression maps showing spatial distribution of CPS1, ARG1, TFF2, S100A6, TOP2A, and nFeature_RNA on the UMAP embedding.

(C) Comparison of clustering and annotation strategies on scVI-corrected UMAP (batch correction via scVI). Left: Leiden clustering (resolution = 0.3). Right: cell state annotation transferred from Seurat, including benign (blue), de_diff_cell (orange), high_diff_cell (green), and prolif_cell (red), showing consistent separation of differentiation states in the scVI embedding.

#### Supplementary Figure 2: Cell state–specific regulatory programs inferred by pySCENIC.

(A) Dot plot showing regulon specificity scores (RSS) across cell states (high_diff_cell, prolif_cell, de_diff_cell, benign). Dot color indicates RSS Z-score, and dot size represents the percentage of cells expressing each regulon.

(B–D) RSS ranking plots for high_diff_cell (B), de_diff_cell (C), and prolif_cell (D). Regulons are ranked by RSS values (x-axis), with corresponding RSS scores on the y-axis; key state-specific regulons are highlighted.

#### Supplementary Figure 3: Altered TAL1 DNA methylation and DNA methyltransferase expression in hepatocellular carcinoma

(A) Box plot showing DNA methylation levels (β values) of TAL1-associated CpG sites in TCGA LIHC tumor (blue) versus normal (red) samples (UCSC Xena). The y-axis represents β values (unitless), and the x-axis indicates individual CpG probes.

(B–C) Box plots showing RNA expression levels [log₂(TPM + 1)] of DNMT3A (B) and DNMT3B (C) in TCGA LIHC tumor (red) versus normal (gray) tissues (GEPIA2). Tumor samples show higher expression of both DNMT3A and DNMT3B (DNMT3A: *p* < 0.01).

#### Supplementary Figure 4: Cellular heterogeneity of CD45⁺ immune cells and T cell subsets.

(A) UMAP visualization of CD45⁺ immune cells colored by annotated cell types.

(B) Dot plot showing expression levels (color intensity) and proportion of expressing cells (dot size) for marker genes across immune cell types defined in (A).

(C) UMAP visualization of T cell subsets colored by subtype annotation.

(D) Dot plot showing expression levels and cell fractions of marker genes across T cell subsets defined in (C).

(E) Feature plots showing the spatial expression of TRAC, TRBC1, KLRB1, TRDC, TRGC1, TRGC2, NKG7, KLRF1, CD3E, CD8A, and CD4 on UMAP embedding, with color intensity indicating expression levels per cell.

#### Supplementary Figure 5: Cross-cohort validation of CD4 T cell heterogeneity and functional states

(A) Feature plots showing expression of CD4 Tfh-associated genes (CXCL13, CD200, TOX2, IL21) on the UMAP space of CD4 T cells. Color intensity indicates mean expression level per cell, highlighting transcriptional features distinguishing Tfh from GZMK CD4 T cells.

(B) Analysis of Liu’s T cell dataset processed in Scanpy with batch correction by scVI. T cells were defined as TCR-positive cells. UMAP shows clustering by the Leiden algorithm (resolution = 0.4), TCR-based annotation, and expression of key markers, indicating that GZMB-expressing populations also include CD4 T cells.

(C) Box plot showing proportions of CD4 T cell subpopulations (CD4_T, GZMB_CD4_T, GZMK_CD4_T, Tfh, Treg) in anti-PD-1 non-responders (NR, light blue) and responders (R, dark blue). Each dot represents an individual patient, with increased Tfh abundance observed in responders.

#### Supplementary Figure 6: Pseudotime trajectory analysis of CD4⁺ T-cell differentiation

(A) Monocle trajectory visualization showing the inferred developmental manifold of CD4⁺ T cells from the Magen dataset. Cells are colored and labeled by cluster identity, with branch structures and terminal states indicated.

(B) Monocle trajectory visualization colored by pseudotime, illustrating the inferred temporal progression across the cell population.

(C) Two-dimensional plots depicting dynamic expression of representative marker genes along pseudotime for distinct differentiation paths. The central curve represents the fitted trend of gene expression across pseudotime, and the shaded region denotes the 95% confidence interval. B_path: from naïve CD4⁺ T cells (marked by CCR7, LEF1 and TCF7) to GZMB⁺ CD4⁺ T cells. K_path: from naïve CD4⁺ T cells to GZMK⁺ CD4⁺ T cells. Self_path: from naïve CD4⁺ T cells to mature CD4⁺ T cells (not assigned to defined subtypes). Tfh_path: from naïve CD4⁺ T cells to Tfh cells. Treg_path: from naïve CD4⁺ T cells to Treg cells.

#### Supplementary Figure 7: Pseudotime trajectory analysis of CD8⁺ T-cell differentiation

(A) Monocle trajectory visualization showing the inferred developmental manifold of CD8⁺ T cells. Cells are colored and labeled by cluster identity, with branch structures and terminal states indicated.

(B) Monocle trajectory colored by pseudotime, illustrating the inferred temporal progression across the cell population.

(C) Two-dimensional plots depicting dynamic expression of representative marker genes along pseudotime for distinct differentiation paths. The central curve represents the fitted trend of gene expression across pseudotime, and the shaded region denotes the 95% confidence interval. GZMB_path: from naïve CD8⁺ T cells (marked by CCR7, LEF1 and TCF7) to GZMB⁺ CD8⁺ T cells. GZMK_path: from naïve CD8⁺ T cells to GZMK⁺ CD8⁺ T cells, and then Tex cells.

(D) Feature plots displaying the expression of CXCR3, CXCR6, CCR5, and CX3CR1 overlaid on the UMAP embedding. Note that CX3CR1 is expressed exclusively on GZMB⁺ CD8⁺ T cells.

#### Supplementary Figure 8: CD8 T Cell subtype–specific regulatory programs inferred by pySCENIC.

(A) Dot plot showing regulon specificity scores (RSS) across cell subtypes (Naïve_CD8_T, GZMB_CD8_T, GZMK_CD8_T, Tex). Dot color indicates RSS Z-score, and dot size represents the percentage of cells expressing each regulon.

(B, C) RSS ranking plots for GZMB_CD8_T (B) and GZMK_CD8_T (C). Regulons are ranked by RSS values (x-axis), with corresponding RSS scores on the y-axis; key state-specific regulons are highlighted.

(D) Dot plot showing the metabolic pathway activity scores at the single-cell level.

#### Supplementary Figure 9: Remodeling of the myeloid compartment in liver tumors.

(A) UMAP visualization of myeloid cell populations in normal (left) and tumor (right) liver tissues with cell-type annotations. Kupffer cells are predominantly enriched in normal liver, whereas two additional tumor-associated subsets, Lipo_Mph_New and Inf_Mono_Mph_New, are expanded in tumor tissues.

(B–D) Feature plots showing the expression patterns of selected marker genes across the myeloid cell population.

(B) SEPP1, EGR1, FOLR2, and CD5L are co-upregulated in Kupffer cells and Lipo_Mph_Propria.

(C) C1QB, FTL, C1QA, and MARCO represent canonical Kupffer cell markers.

(D) KIT, AREG and GATA3 mark ILCs, while KIT and MS4A2 marks mast cells.

#### Supplementary Figure 10: B-cell spatial distribution, cellular sources of CXCL12, and transcriptional features of B-cell subsets in normal liver and HCC.

(A) UMAP visualizations of B cell populations in normal (left) and tumor (right) liver tissues with annotation. Note that plasmablasts are predominantly enriched in normal liver, while the transcriptional profile of atypical memory B cells shows significant shifts in the tumor microenvironment.

(B) UMAP of myeloid cells with annotation (left) and CXCL12 expression (right) overlaid, highlighting expression in Kupffer cells and Lipo_Mph.

(C) UMAP of fibroblast populations (left) and corresponding CXCL12 expression (right), showing enrichment in stellate cells.

(D) Feature plots of representative up- and down-regulated genes in the C7 population. C7 upregulates cytoskeletal regulators (ACTB, ACTG1, COTL1, RAC2) and downregulates key homing receptors (CCR7, CXCR4, CXCR5, GPR183), consistent with a highly migratory phenotype.

(E) Feature plots of representative stress-response genes (DNAJB1, HSPA1A, HSP90AA1, HSPE1) upregulated in the C8 population.

(F) Feature plots showing CXCR3, PRDM1, IRF4, and TNFRSF17 expression mapped onto the UMAP embedding. The C6 subset expresses PRDM1 and IRF4 but shows lower TNFRSF17 expression.
